# Evidence for S_331_-G-S-L within the amyloid core of myocilin olfactomedin domain fibrils based on low-resolution 3D solid-state NMR spectra

**DOI:** 10.1101/2024.08.09.606901

**Authors:** Emily G. Saccuzzo, Alicia S. Robang, Yuan Gao, Bo Chen, Raquel L. Lieberman, Anant K. Paravastu

## Abstract

Myocilin-associated glaucoma is a protein-conformational disorder associated with formation of a toxic amyloid-like aggregate. Numerous destabilizing single point variants, distributed across the myocilin olfactomedin β-propeller (OLF, myocilin residues 245-504, 30 kDa) are associated with accelerated disease progression. *In vitro*, wild type (WT) OLF can be promoted to form thioflavin T (ThT)-positive fibrils under mildly destabilizing (37°C, pH 7.2) conditions. Consistent with the notion that only a small number of residues within a protein are responsible for amyloid formation, 3D ^13^C-^13^C solid-state NMR spectra show that OLF fibrils are likely to be composed of only about one third of the overall sequence. Here, we probe the residue composition of fibrils formed *de novo* from purified full-length OLF. We were able to make sequential assignments consistent with the sequence S_331_-G-S-L_334_. This sequence appears once within a previously identified amyloid-prone region (P1, G_326_AVVYSGSLYFQ) internal to OLF. Since nearly half of the pairs of adjacent residues (di-peptides) in OLF occur only once in the primary structure and almost all the 3-residue sequences (tri-peptides) are unique, remarkably few sequential assignments are necessary to uniquely identify specific regions of the amyloid core. This assignment approach could be applied to other systems to expand our molecular comprehension of how folded proteins undergo fibrillization.

**TOC GRAPHIC:** 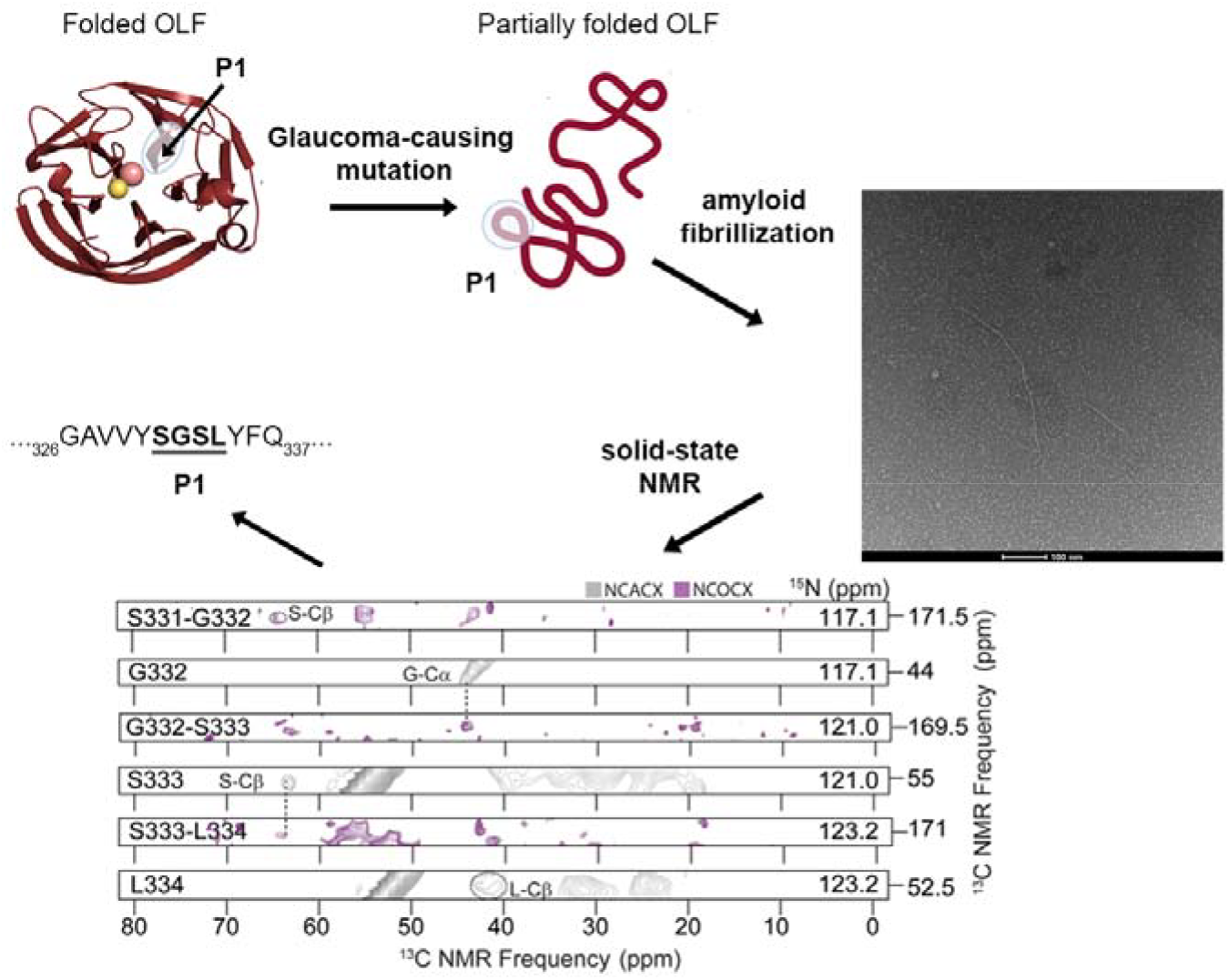

---

A decade ago, the 30 kDa olfactomedin (OLF) domain of myocilin was added to the list of disease-associated proteins that can aggregate to form fibrils with characteristics of amyloid^1^. Numerous missense mutations within OLF are genetically associated with ∼5% of cases of primary open angle glaucoma (POAG), leading to early-onset vision loss^2, 3^. Myocilin is robustly expressed in the trabecular meshwork, an anatomical region of the anterior eye that loses cellularity over time and is diseased in most forms of glaucoma^4^. Whereas full-length myocilin is a secreted glycoprotein^5^, mutant myocilins aggregate within the endoplasmic reticulum of cells^6, 7^ causing cell stress and cell death^7^. Compared to wild-type (WT) OLF, disease variants are thermally destabilized^8^ and adopt non-native tertiary states that facilitate aggregation at pH 7.2 and 37 °C^9^.

At present, there are limited studies involving amyloid formation by large, non-model proteins such as OLF, a 30 kDa 5-bladed β-propeller with an internal metal center (Figure 1a). The cross-β motif common to amyloid structure is compatible with a wide range of peptide and protein sizes, including peptides as small as 100 Da (around two amino acids)^10-12^ and proteins as large as 95 kDa (thousands of amino acids)^13^. Previous results for larger proteins, such as transthyretin^14^ or β-2-microglobulin^15^, indicate that the amyloid core involves only a portion of the residues. Other residues retain partially folded structures, constituting turn regions between β-strands in the amyloid structure, or remain unstructured. The mechanism of fibril formation is thought to proceed by the templating of amyloid prone region(s) exposed during partial unfolding until mature fibrils precipitate out of solution^16^. Model proteins have been shown to adopt partially folded structures upon chemical denaturation^17, 18^, addition of heat^19, 20^, mutation^16^, dissociation of a higher ordered oligomer such as tetramer^14^ or dimer^21^ to monomer, or loss of metal^22^. Overall, we are still lacking a fundamental understanding of how partial unfolding is initiated and propagated for fibril formation and how many amino acids in a large protein (> 20 kDa) are involved in forming fibril cores.

**Figure 1.**
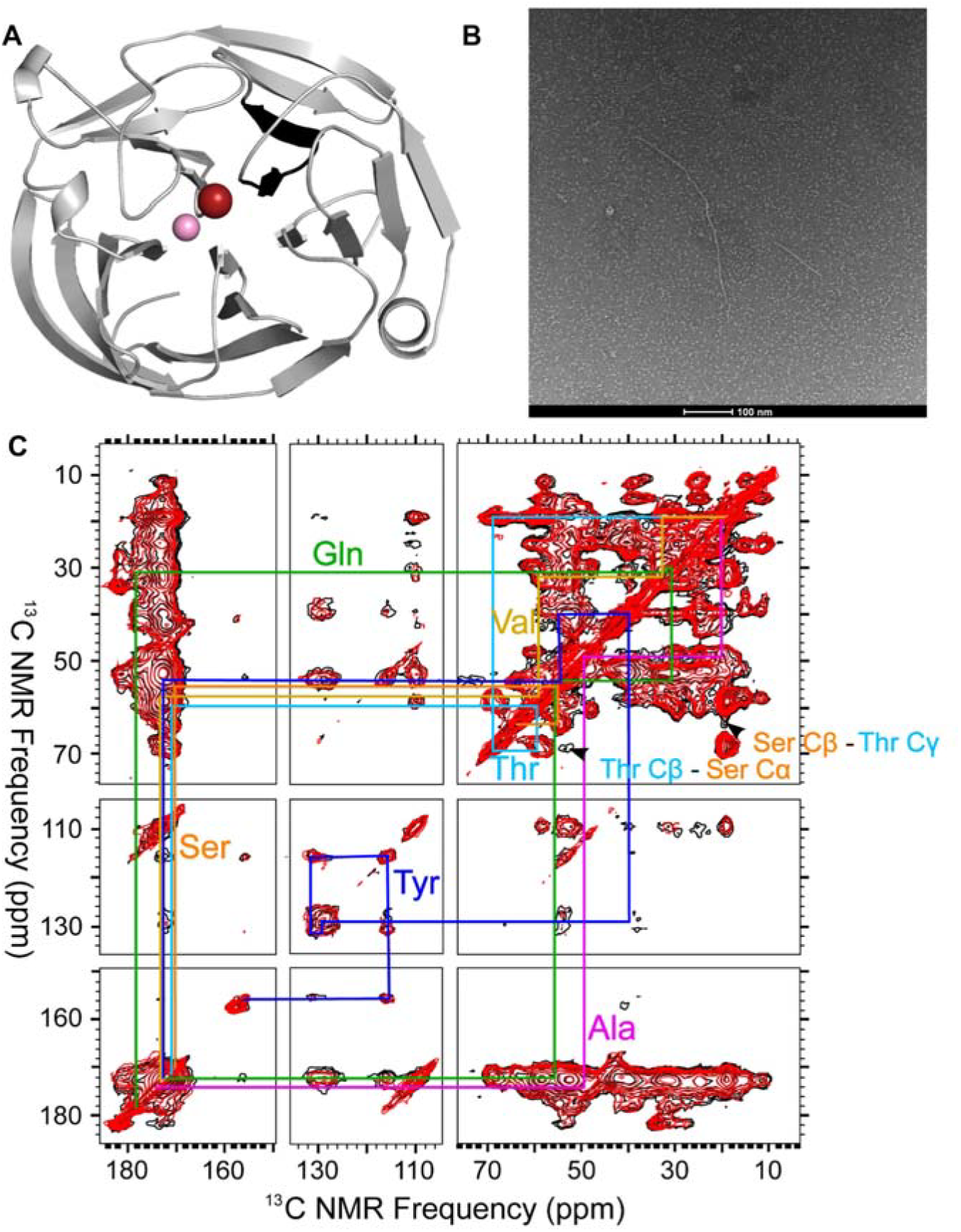
2D ^13^C-^13^C solid-state NMR and TEM of OLF fibrils. (A) Location of P1 peptide identified in black on the OLF structure (PDB 4WXQ). The internal calcium and sodium ions are represented as pink and red spheres, respectively. (B) Representative negative stain TEM of an OLF fibril deposited on a positively charged grid surface, fibril is stained with 2% uranyl acetate (UA) and image was collected on a 120 kV Talos electron microscope. (C) 2D-DARR spectrum. Cross-peaks from intra-residue ^13^C-^13^C correlations shown in colored pathways or labeled with arrows. Comparing red spectrum (50 ms DARR mixing) to black spectrum (100 ms DARR mixing) identifies adjacent residues.

Here we employed solid-state NMR to identify approximately how many, as well as what specific residues, participate in the amyloid core of the full-length myocilin WT OLF formed *de novo* e.g. without seeding. Further, we sought experimental evidence of the previously reported amyloid-forming stretch of residues 326 to 337 within the OLF fibril core. This residue stretch corresponds to peptide P1 (G_326_AVVYSGSLYFQ, Figure 1a), which was identified by amyloid prediction software and experimentally shown to form amyloid-like fibrils^9^. We previously reported a possible connection between peptide P1 and OLF protein aggregation: AFM showed similarities in nanoscale morphologies between P1 amyloid and OLF fibrils formed under different protocols: OLF fibrils formed via incubation at 37°C with gentle rocking appear morphologically similar to those formed by P1 peptide alone^9^. Figure 1b shows an example of fibril morphology of OLF fibrils observed from negative stain transmission electron microscopy (TEM). For the NMR experiments herein, we used previously established protocols to express and purify uniformly ^13^C and ^15^N labeled OLF^23^ and form fibrils previously shown to exhibit multiple characteristics of amyloid^9^.

Initial two-dimensional (2D) ^13^C-^13^C dipolar assisted rotational resonance (DARR) spectra of OLF fibril samples identified residue types involved in forming the aggregated species. To determine the residues involved in forming the amyloid core, we first collected 2D DARR at 50 ms mixing time (Figure 1c, red) to partially assign ^13^C chemical shifts to ^13^C atoms within a single residue. Figure 1c shows preliminary chemical shift assignments consistent with I, G, P, M, K, L, F, Y, R, and D residues, based on known chemical shift ranges from the Biological Magnetic Resonance Data Bank^24, 25^. We observed ^13^C NMR peak linewidths in the range of ∼ 2 to 3 ppm, which are more broad than previously reported linewidths of 1 to 2 ppm within highly ordered Alzheimer’s amyloid-β fibrils^26, 27^. However, linewidths of 2 to 3 ppm are not atypical for amyloid formation and can indicate a degree of heterogeneity in the structure.

We conducted additional 2D DARR measurements on the fibrils at 100 ms mixing time to observe off-diagonal crosspeaks between adjacent residues (Figure 1c, black). We detected inter-residue crosspeaks between S and T residues (Figure 1c). Since the appearance of 2D DARR crosspeaks does not provide information about the sequential arrangement or the N-to C-terminus direction of the residues, we also analyzed the OLF sequence for direction-dependent S-T and T-S connections. Analysis of the OLF amino acid sequence revealed three S-T connections, one of which directly precedes the previously identified P1 peptide sequence (Supp. Fig S1a-b).

To determine whether stretches of residues from the P1 peptide comprise the fibril core of OLF, we conducted 3D NMR NCACX and NCOCX experiments. In these experiments, spin polarization is transferred first from proton to the backbone ^15^N site, and then to ^13^C sites (^13^Cα for NCA transfer and ^13^C’ for NCO transfer) using optimized adiabatic cross polarization to make sequential backbone chemical shift assignments. These transfers were then extended with a non-specific ^13^C-homonuclear transfer step obtained with DARR mixing to obtain “CX,” where X denotes any carbon. We looked for serine and threonine residues within the NCACX and NCOCX spectra collected on the OLF fibrils^28^, as these residues have distinguishable chemical shifts^24, 25^. Based on Gaussian peak fitting to ^15^N-^13^C 2D projections, we estimated 7 S peaks and 10 T peaks involved in the formation of the fibril (Supp Fig. S2). Our estimation represents, on average, about one third of the amount of each residue in the full OLF sequence (Supp. Table S1), suggesting ∼ 36% of the 277 amino acids in the OLF sequence (∼ 98 residues) are involved in forming the fibril core. However, overlapping peaks are possible and less than 98 residues may be involved in forming the fibril core.

The high number of di-peptide and tri-peptide stretches that only appear once within the OLF sequence (124 unique di-peptides, 267 unique tri-peptides) (Supp Fig. S3) allowed for unambiguous assignment of a unique 4-peptide stretch in the OLF fibril spectra. The N/CX and C/CX strip plots, or strips, illustrate the assignment procedure (Figure 2). Both spectra were first scanned at ^15^N frequencies around 120 ppm (the typical range for ^15^N backbone atoms) for unique fingerprint patterns belonging to either G or S^24, 25^. Namely, G only has two peaks in the carbon dimension for any given nitrogen frequency (Cα ∼ 41 ppm, C’ ∼ 171 ppm), and S has a unique Cα/Cβ crosspeak at ∼ 54 ppm/ ∼ 62 ppm^29^. We identified a G spin system at a ^15^N frequency of 121.0 ppm in a strip from the NCOCX experiment (Figure 2a). To complete a sequential assignment for the S residue in the *i* + 1 direction the corresponding C/CX strip was scanned at 121.0 ppm, and an S Cα/Cβ crosspeak (55.4 ppm/ 63.4 ppm) was detected (Figure 2a). These initial chemical shift assignments established the sequential connection of G-S. Although there are 23 G residues within the OLF sequence, the sequential G-S dipeptide is present only once and is within the previously identified amyloid-prone region of P1 (Supp Fig. S1). Thus, G_332_-S_333_ was assigned with confidence.

**Figure 2.**
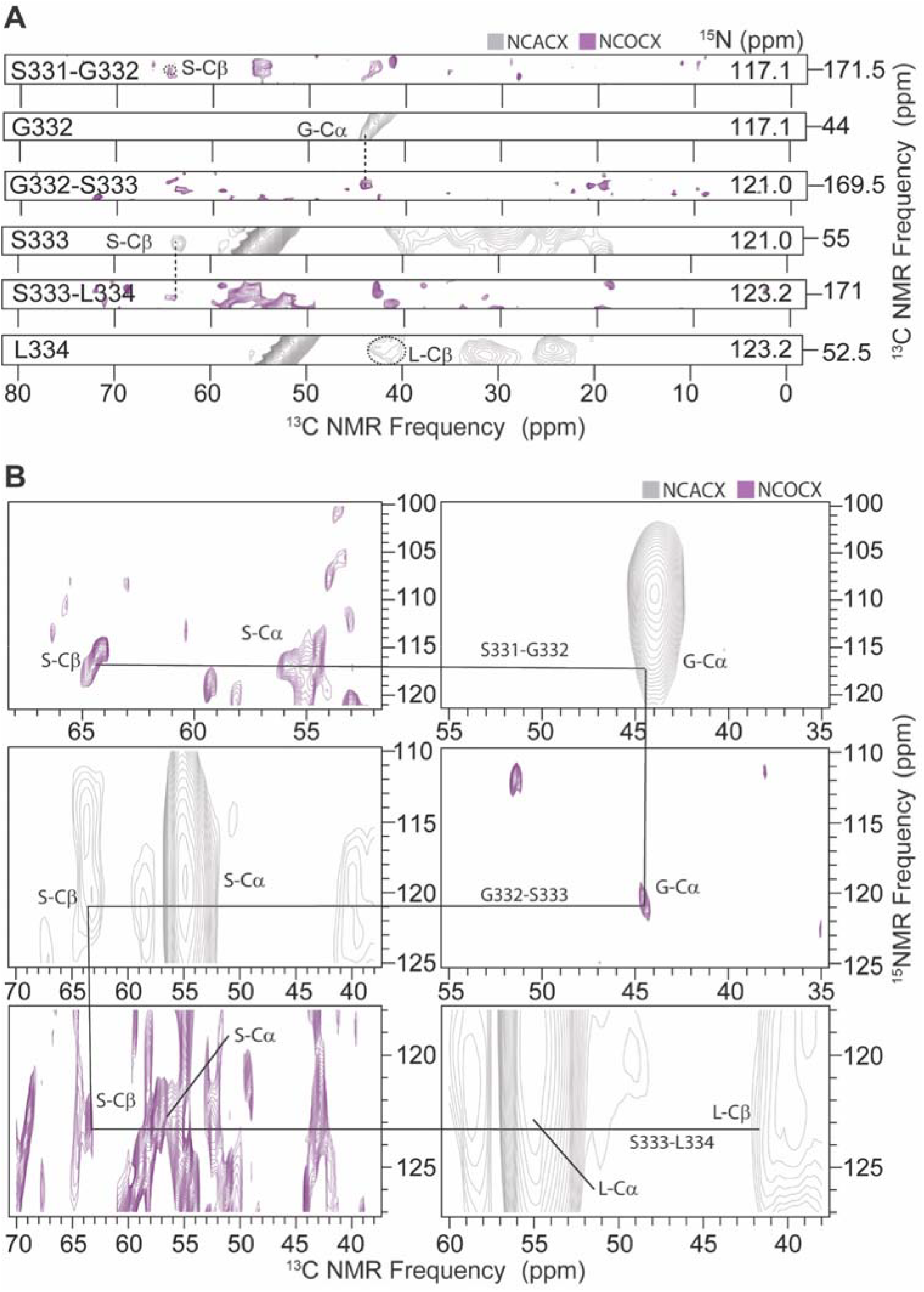
Assignment of P1 sequence “S_331_-G-S-L_334_” in OLF fibrils. **(A)** C/Cx strip plots were extracted from NCACX and NCOCX 3D spectra. Residues were connected via ^15^N chemical shifts and identified by their characteristic ^13^C chemical shifts (labeled). **(B)** Due to relatively poor spectral resolution in the nitrogen dimension, assignments were predominantly made using C/Cx strip plots. N/Cx strip plots were used for validation of these assignments. Alternating N/Cx strips from the NCOCX and NCACX spectra are shown for each residue, and lines are drawn in to show sequential connections. Chemical shifts for each amino acid are listed in Figure 3 and Table S1.

To support the G_332_-S_333_ sequential assignment, we next conducted a series of sequential walks through the set of 3D spectra. If assigned correctly, G_332_-S_333_ should be flanked by the S_331_residue in the *i*-1 direction, (Figure 3, Supp. Fig. S1). Thus, we looked for signals that support the connection for S_331_-G_332_. We scanned C/CX strips at ^15^N frequencies to locate possible S Cα/Cβ crosspeaks. At a ^15^N frequency of 117.1 ppm, we observed both a peak at 44.5 ppm, which is the G_332_ Cα frequency observed in the NCOCX spectrum, and a S Cα/Cβ crosspeak (55.9 ppm/ 64.2 ppm) (Figure 2b). These sequential walks support the assignment of “S_331_-G-S_333_”. Thus far, the unique chemical shifts of S Cα and Cβ allowed chemical shift assignments from NCACX and NCOCX spectra that support S connections to either side of G. These assignments illustrate that we can identify sequential residues even from low-resolution 3D spectra.

**Figure 3.**
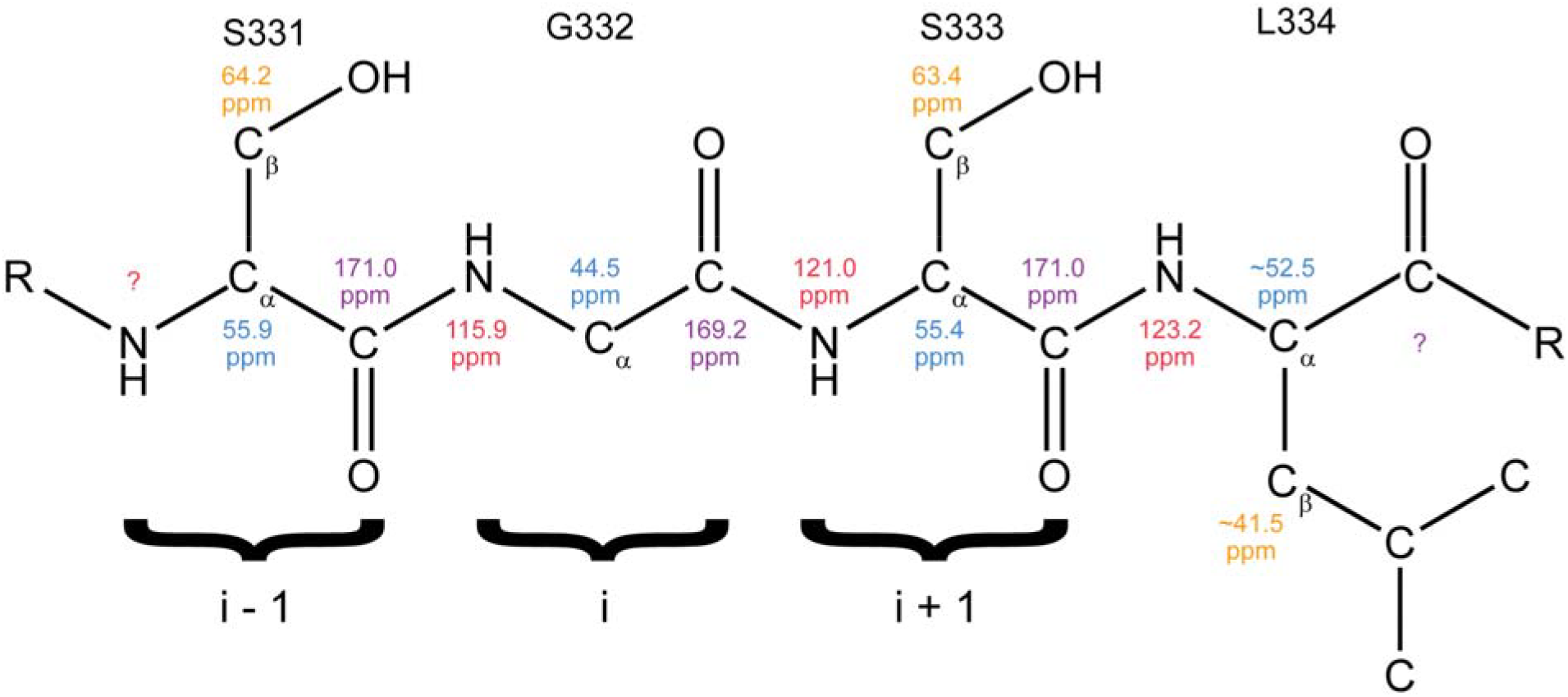
Assigned S_331_-G-S-L_334_ peptide with corresponding chemical shifts. Representation of the assigned stretch S_331_-G-S-L_334_, with chemical shifts for Cα in blue, Cβ in orange, CO in purple and N in red. An example of *i* −1, *i*, and *i* + 1 directionality, which is used in the text, is labeled.

From analysis of the OLF sequence, we observed that S_331_-G-S_333_, like the initial G_332_-S_333_, is a unique peptide stretch in the OLF sequence and the only possible assignment in the *i* + 1 direction is L_334_ (Figure 3, Supp. Fig. S1). Unfortunately, uniquely identifying L peaks is expected to be more challenging because other amino acids have overlapping ^13^C chemical shifts^29^. The identification of a Cα/Cβ crosspeak at ∼ 51 ppm/ ∼ 41 ppm would be consistent with an L residue. Nonetheless, we were able to locate the S_333_ Cα/Cβ crosspeak (55.4 ppm/ 63.4 ppm) in the NCOCX spectrum, at a ^15^N frequency of 123.2 ppm. This nitrogen frequency was then scanned in the NCACX spectrum, and a C/CX crosspeak at ∼ 52.5 ppm/ ∼ 41.5 ppm was identified, which could be attributed to L_334_. While it is difficult to confirm that these peaks are uniquely due to L_334_ because of the low resolution, the spectra do not contradict an S_331_-G-S-L_334_ assignment. Figure 3 and Supplementary Table S2 show the full set of current chemical shift assignments for carbon and nitrogen atoms within S_331_-G-S-L_334_ residues.

Either ends of S_331_-G-S-L_334_ are capped by Y residues, which is another difficult chemical shift assignment in multidimensional NMR. Although Y has unique chemical shifts in the aromatic region, aromatic peaks typically have minimal signal to noise and are difficult to detect in multidimensional NMR^30^. Our 2D DARR spectra displayed possible ^13^C peaks for Y (Figure 1b). However, we were not able to detect any well-resolved peaks in the 3D spectra connecting Y to S residues. While we cannot make connections beyond S_331_-G-S-L_334_, we are confident in this assignment from our analysis of the OLF sequence, which revealed this tetrapeptide stretch is unique.

Building on indirect methods that suggested P1 as an amyloid-prone region in OLF, we now have evidence that the amyloid core formed by OLF fibrils comprises at least in part by the sequence represented by P1^9^. We identified crosspeaks from NCACX and NCOCX measurements consistent with S_331_-G-S-L_334_ connections and determined that this tetra-peptide stretch is unique in the entire OLF sequence. In natively folded OLF, P1 forms a β-hairpin and is buried in the interior of the protein (Figure 1a). Thus, like better studied model systems^14, 15^, OLF must undergo partial unfolding to expose aggregation-prone region(s) for fibrillization.

Further, our results demonstrate that solid-state NMR techniques can be adapted for the identification of residues in the amyloid core of samples that give relatively low-resolution spectra. Rather than assigning all peaks in a spectrum to determine a molecular structure, we combined low-resolution spectra with knowledge of the amino acid sequence to determine whether a predicted stretch of residues participates in the amyloid core. High resolution spectra can be obtained by increasing homogeneity, for example by seeding^31^, to allow a single morphology to predominate. However, it is now thought that heterogeneous fibrils may be disease relevant^32^. The NMR measurements presented in this work alone are not sufficient to comment on the nature of the heterogeneity within fibrils formed by WT OLF and there is no biological evidence to suggest that pathologically relevant fibrils formed by mutant myocilin are highly homogeneous. Nevertheless, our study shows that in spite of the challenges associated with heterogeneous fibril samples, even limited assignments from low-resolution spectra can provide novel insight into the process of fibrillization of complex systems.

## Experimental Methods

### Expression and purification of ^13^C/^15^N labeled OLF

The plasmid for WT OLF, which encodes for an N-terminal maltose binding protein (MBP) and Tobacco Etch Virus protease (TEV) cleavage site before the OLF domain^33^ was used for expression in *E. coli* T7 Express cells using the protocol previously reported^34^, with minor adjustments. Prior to resuspension in 500 mL minimal media, the T7 express cells (pelleted from 500 mL of rich media) were washed twice with ∼100 mL of 1× M9 salts (26 g·L^−1^ Na_2_HPO_4_·7H_2_O, 3 g·L^−1^ KH_2_PO_4_, 0.5 g·L^1^ NaCl, 1 g·L^−1 15^NH_4_Cl). 20 mL of 20% ^13^C glucose per liter of media (sterile filtered) was used as the carbon source. The minimal media was also supplemented with 1 mL of 1000X trace-elements solution (40.8 mM CaCl_2_*2H_2_O, 21.6 mM FeSO_4_*7H_2_O, 6.1 mM MnCl_2_*4H_2_O, 3.4 mM CoCl_2_*6H_2_O, 2.4 mM ZnSO_4_*7H_2_O, 1.8 mM CuCl_2_*2H_2_O, 0.3 mM Boric Acid, 0.2 mM ammonium molybdate, 5 g/L EDTA) (sterile filtered), 0.1 mL of 1 M CaCl_2_ (sterile filtered) and 2.5 mL BME vitamins (Sigma Aldrich, B6891−100ML) (sterile)). Cells were lysed and OLF was purified as previously described ^33^. Sample purity was assessed with standard SDS-PAGE with Coomassie staining.

### OLF fibrillization

Uniformly ^13^C/^15^N labeled WT OLF in 10 mM Na/K phosphate buffer with 0.2 M NaCl pH 6.8 was concentrated to 30 μM using Amicon filtration devices with a 10 kDa MWCO. Absorbance at 280 nm was measured on a UV/Vis spectrometer and concentration was calculated using the Beer-Lambert law with an extinction coefficient of 68425 M^−1^cm^−1^. Purified ^13^C/^15^N labeled OLF (30 μM, 2 mL) was added to a 5 mL microtube (Cat. No. T2076S-CA). The tubes were sealed with parafilm and added to a 2D rocker in a 37°C incubator for 96 hours. The experiment was conducted in triplicate. The same procedure was performed with unlabeled WT OLF to generate fibrils for TEM imaging.

### TEM imaging

Microscopy was conducted at the Emory University Robert P. Apkarian Integrated Electron Microscopy Core. Fibrils (2 μL) were deposited directly onto a positively glow-discharged, 400 mesh carbon-coated copper grid. After a 1-minute incubation, the sample was blotted from the grid with Whatman no. 4 filter paper and washed 2x with water droplets. The grids were negatively stained with a drop of 2% uranyl acetate and imaged using a FEI Talos 120 kV standard transmission electron microscope equipped with a LaB6 and 4k Ceta detector.

### 2D and 3D solid-state NMR experiments

All solid-state NMR experiments were conducted on a Bruker narrow-bore 11.7 T magnet (^1^H frequency of 500 MHz), equipped with a 3.2 mm low-E HCN magic angle spinning (MAS) probe. Samples were ultracentrifuged for 30 minutes into Bruker 3.2 mm NMR rotors at 4 °C and 150,000 xg using a Beckman Optima XPN−100 centrifuge with Ultra-clear tubes in a SW-41 Ti swinging bucket rotor. Rotors were paced using a custom-made funnel widget to channel the centrifuge pellet into the NMR rotor.

For 2D ^13^C-^13^C DARR experiments, continuous ^1^H irradiation was used with a power corresponding to 11 kHz during the ^13^C-^13^C mixing period, which is the same as the MAS spinning rate ^35^. Two-pulse-phase modulation (TPPM) was the proton decoupling method used with a power set to 110 kHz. The pulse power was set to 50 kHz for all ^13^C pulses, including the cross-polarization spin lock and the hard pulses. The mixing period was 50 ms for detecting inter-residue crosspeaks and 100 ms for detecting intra-residue crosspeaks. Signal averaging was conducted over 36 h.

For sequence-specific assignments, NCOCX/NCACX spectra were acquired. Frequency-selective heteronuclear ^15^N-^13^C correlations, specifically between amide ^15^N and the ^13^Cα (NCA transfer) or the C’ (NCO transfer), were done using optimized adiabatic cross polarization^28^. The pulse sequence hNCaCX3D.dcp was used for both NCACX and NCOCX experiments. For the first step of ^1^H to ^15^N polarization, we used the following pulse powers and lengths: 4 ms duration, 50 kHz ^15^N pulse power, 50 to 100% ^1^H pulse power ramp to 100 kHz. For the second step of ^15^N to ^13^C polarization, we used the following pulse powers and lengths: 9 ms duration, 45 kHz ^15^N pulse power optimized for the best ^15^N to ^13^C transfer, 100 kHz ^1^H decoupling power, 71 kHz ^13^C pulse power with 50% tangential shape. The ^15^N to ^13^Cα transfers were then extended with a non-specific ^13^C-homonuclear transfer step done with a DARR mixing time of 30 ms to obtain “CX,” which denotes any carbon. Signals for NCACX were averaged over 3 days. Signals for NCOCX were averaged over 6 days. Appropriate 2D slices were extracted from both the NCOCX and NCACX spectra and exported into Mathematica.

## Associated Content

Supporting Figures S1-S3 and Tables S1-S2 are available in the accompanying Supporting Information file.

## Author Information

Correspondence to Anant K. Paravastu or Raquel L. Lieberman.

## Acknowledgements and funding

This work was supported by NIH grants R01EY021205 and RF1AG073434. EGS was supported by 5T32EY007092 and GAANN P200A210014. We acknowledge the following core facilities: Georgia Tech Petit Institute for Bioengineering and Biosciences and NMR Center, and the Emory University Robert P. Apkarian Integrated Electron Microscopy Core (IEMC). IEMC is subsidized by the School of Medicine and Emory College of Arts and Sciences and Georgia Clinical & Translational Science Alliance of the National Institutes of Health under award number UL1TR000454. We thank Dr. Dustin Huard and Hailee Scelsi for valuable discussions and critical review of the manuscript.

## References

(1) Orwig, S. D.; Perry, C. W.; Kim, L. Y.; Turnage, K. C.; Zhang, R.; Vollrath, D.; Schmidt-Krey, I.; Lieberman, R. L. Amyloid fibril formation by the glaucoma-associated olfactomedin domain of myocilin. J Mol Biol 2012, 421 (2-3), 242–255. DOI: 10.1016/j.jmb.2011.12.016.

(2) Kwon, Y. H.; Fingert, J. H.; Kuehn, M. H.; Alward, W. L. Primary open-angle glaucoma. N Engl J Med 2009, 360 (11), 1113–1124. DOI: 10.1056/NEJMra0804630.

(3) Fingert, J. H.; Heon, E.; Liebmann, J. M.; Yamamoto, T.; Craig, J. E.; Rait, J.; Kawase, K.; Hoh, S. T.; Buys, Y. M.; Dickinson, J.; et al. Analysis of myocilin mutations in 1703 glaucoma patients from five different populations. Hum Mol Genet 1999, 8 (5), 899–905. DOI: 10.1093/hmg/8.5.899.

(4) Alvarado, J.; Murphy, C.; Juster, R. Trabecular meshwork cellularity in primary open-angle glaucoma and nonglaucomatous normals. Ophthalmology 1984, 91 (6), 564–579. DOI: 10.1016/s0161-6420(84)34248-8 From NLM Medline.

(5) Tamm, E. R. Myocilin and glaucoma: facts and ideas. Prog Retin Eye Res 2002, 21 (4), 395–428. DOI: 10.1016/s1350-9462(02)00010−1.

(6) Joe, M. K.; Sohn, S.; Hur, W.; Moon, Y.; Choi, Y. R.; Kee, C. Accumulation of mutant myocilins in ER leads to ER stress and potential cytotoxicity in human trabecular meshwork cells. Biochem Biophys Res Commun 2003, 312 (3), 592–600. DOI: 10.1016/j.bbrc.2003.10.162.

(7) Yam, G. H.; Gaplovska-Kysela, K.; Zuber, C.; Roth, J. Aggregated myocilin induces russell bodies and causes apoptosis: implications for the pathogenesis of myocilin-caused primary open-angle glaucoma. Am J Pathol 2007, 170 (1), 100–109. DOI: 10.2353/ajpath.2007.060806.

(8) Burns, J. N.; Turnage, K. C.; Walker, C. A.; Lieberman, R. L. The stability of myocilin olfactomedin domain variants provides new insight into glaucoma as a protein misfolding disorder. Biochemistry 2011, 50 (26), 5824–5833. DOI: 10.1021/bi200231x.

(9) Hill, S. E.; Donegan, R. K.; Lieberman, R. L. The glaucoma-associated olfactomedin domain of myocilin forms polymorphic fibrils that are constrained by partial unfolding and peptide sequence. J Mol Biol 2014, 426 (4), 921–935. DOI: 10.1016/j.jmb.2013.12.002.

(10) de Groot, N. S.; Parella, T.; Aviles, F. X.; Vendrell, J.; Ventura, S. Ile-phe dipeptide self-assembly: clues to amyloid formation. Biophys J 2007, 92 (5), 1732–1741. DOI: 10.1529/biophysj.106.096677.

(11) Ji, W.; Yuan, C.; Zilberzwige-Tal, S.; Xing, R.; Chakraborty, P.; Tao, K.; Gilead, S.; Yan, X.; Gazit, E. Metal-Ion Modulated Structural Transformation of Amyloid-Like Dipeptide Supramolecular Self-Assembly. ACS Nano 2019, 13 (6), 7300–7309. DOI: 10.1021/acsnano.9b03444.

(12) Ji, W.; Yuan, C.; Chakraborty, P.; Makam, P.; Bera, S.; Rencus-Lazar, S.; Li, J.; Yan, X.; Gazit, E. Coassembly-Induced Transformation of Dipeptide Amyloid-Like Structures into Stimuli-Responsive Supramolecular Materials. ACS Nano 2020, 14 (6), 7181–7190. DOI: 10.1021/acsnano.0c02138.

(13) Solomon, J. P.; Page, L. J.; Balch, W. E.; Kelly, J. W. Gelsolin amyloidosis: genetics, biochemistry, pathology and possible strategies for therapeutic intervention. Crit Rev Biochem Mol Biol 2012, 47 (3), 282–296. DOI: 10.3109/10409238.2012.661401.

(14) Yee, A. W.; Aldeghi, M.; Blakeley, M. P.; Ostermann, A.; Mas, P. J.; Moulin, M.; de Sanctis, D.; Bowler, M. W.; Mueller-Dieckmann, C.; Mitchell, E. P.; et al. A molecular mechanism for transthyretin amyloidogenesis. Nat Commun 2019, 10 (1), 925. DOI: 10.1038/s41467-019-08609-z.

(15) Smith, D. P.; Jones, S.; Serpell, L. C.; Sunde, M.; Radford, S. E. A systematic investigation into the effect of protein destabilisation on beta 2-microglobulin amyloid formation. J Mol Biol 2003, 330 (5), 943–954. DOI: 10.1016/s0022-2836(03)00687-9.

(16) Chiti, F.; Dobson, C. M. Amyloid formation by globular proteins under native conditions. Nat Chem Biol 2009, 5 (1), 15–22. DOI: 10.1038/nchembio.131.

(17) Vernaglia, B. A.; Huang, J.; Clark, E. D. Guanidine hydrochloride can induce amyloid fibril formation from hen egg-white lysozyme. Biomacromolecules 2004, 5 (4), 1362–1370. DOI: 10.1021/bm0498979.

(18) Goda, S.; Takano, K.; Yamagata, Y.; Nagata, R.; Akutsu, H.; Maki, S.; Namba, K.; Yutani, K. Amyloid protofilament formation of hen egg lysozyme in highly concentrated ethanol solution. Protein Sci 2000, 9 (2), 369–375. DOI: 10.1110/ps.9.2.369.

(19) Arnaudov, L. N.; de Vries, R. Thermally induced fibrillar aggregation of hen egg white lysozyme. Biophys J 2005, 88 (1), 515–526. DOI: 10.1529/biophysj.104.048819.

(20) Ow, S. Y.; Dunstan, D. E. The effect of concentration, temperature and stirring on hen egg white lysozyme amyloid formation. Soft Matter 2013, 9 (40), 9692–9701. DOI: 10.1039/c3sm51671g.

(21) Qin, Z.; Hu, D.; Zhu, M.; Fink, A. L. Structural characterization of the partially folded intermediates of an immunoglobulin light chain leading to amyloid fibrillation and amorphous aggregation. Biochemistry 2007, 46 (11), 3521–3531. DOI: 10.1021/bi061716v.

(22) Teilum, K.; Smith, M. H.; Schulz, E.; Christensen, L. C.; Solomentsev, G.; Oliveberg, M.; Akke, M. Transient structural distortion of metal-free Cu/Zn superoxide dismutase triggers aberrant oligomerization. Proc Natl Acad Sci U S A 2009, 106 (43), 18273–18278. DOI: 10.1073/pnas.0907387106.

(23) Saccuzzo, E. G.; Mebrat, M. D.; Scelsi, H. F.; Kim, M.; Ma, M. T.; Su, X.; Hill, S. E.; Rheaume, E.; Li, R.; Torres, M. P.; et al. Competition between inside-out unfolding and pathogenic aggregation in an amyloid-forming beta-propeller. Nat Commun 2024, 15 (1), 155. DOI: 10.1038/s41467-023-44479-2 From NLM Medline.

(24) Romero, P. R.; Kobayashi, N.; Wedell, J. R.; Baskaran, K.; Iwata, T.; Yokochi, M.; Maziuk, D.; Yao, H.; Fujiwara, T.; Kurusu, G.; et al. BioMagResBank (BMRB) as a Resource for Structural Biology. Methods Mol Biol 2020, 2112, 187–218. DOI: 10.1007/978−1-0716-0270-6_14 From NLM Medline.

(25) Ulrich, E. L.; Akutsu, H.; Doreleijers, J. F.; Harano, Y.; Ioannidis, Y. E.; Lin, J.; Livny, M.; Mading, S.; Maziuk, D.; Miller, Z.; et al. BioMagResBank. Nucleic Acids Res 2008, 36 (Database issue), D402–408. DOI: 10.1093/nar/gkm957 From NLM Medline.

(26) Zech, S. G.; Wand, A. J.; McDermott, A. E. Protein structure determination by high-resolution solid-state NMR spectroscopy: application to microcrystalline ubiquitin. J Am Chem Soc 2005, 127 (24), 8618–8626. DOI: 10.1021/ja0503128 From NLM Medline.

(27) Colvin, M. T.; Silvers, R.; Frohm, B.; Su, Y.; Linse, S.; Griffin, R. G. High resolution structural characterization of Abeta42 amyloid fibrils by magic angle spinning NMR. J Am Chem Soc 2015, 137 (23), 7509–7518. DOI: 10.1021/jacs.5b03997 From NLM Medline.

(28) Pauli, J.; Baldus, M.; van Rossum, B.; de Groot, H.; Oschkinat, H. Backbone and side-chain 13C and 15N signal assignments of the alpha-spectrin SH3 domain by magic angle spinning solid-state NMR at 17.6 Tesla. Chembiochem 2001, 2 (4), 272–281. DOI: 10.1002/1439-7633(20010401)2:4<272::AID-CBIC272>3.0.CO;2-2 From NLM Medline.

(29) Hoch, J. C.; Baskaran, K.; Burr, H.; Chin, J.; Eghbalnia, H. R.; Fujiwara, T.; Gryk, M. R.; Iwata, T.; Kojima, C.; Kurisu, G.; et al. Biological Magnetic Resonance Data Bank. Nucleic Acids Res 2023, 51 (D1), D368–D376. DOI: 10.1093/nar/gkac1050.

(30) Reif, B.; Ashbrook, S. E.; Emsley, L.; Hong, M. Solid-state NMR spectroscopy. Nat Rev Methods Primers 2021, 1. DOI: 10.1038/s43586-020-00002−1 From NLM PubMed-not-MEDLINE.

(31) Paravastu, A. K.; Qahwash, I.; Leapman, R. D.; Meredith, S. C.; Tycko, R. Seeded growth of beta-amyloid fibrils from Alzheimer’s brain-derived fibrils produces a distinct fibril structure. Proc Natl Acad Sci U S A 2009, 106 (18), 7443–7448. DOI: 10.1073/pnas.0812033106.

(32) Sokratian, A.; Ziaee, J.; Kelly, K.; Chang, A.; Bryant, N.; Wang, S.; Xu, E.; Li, J. Y.; Wang, S. H.; Ervin, J.; et al. Heterogeneity in alpha-synuclein fibril activity correlates to disease phenotypes in Lewy body dementia. Acta Neuropathol 2021, 141 (4), 547–564. DOI: 10.1007/s00401-021-02288−1.

(33) Hill, S. E.; Kwon, M. S.; Martin, M. D.; Suntharalingam, A.; Hazel, A.; Dickey, C. A.; Gumbart, J. C.; Lieberman, R. L. Stable calcium-free myocilin olfactomedin domain variants reveal challenges in differentiating between benign and glaucoma-causing mutations. J Biol Chem 2019, 294 (34), 12717–12728. DOI: 10.1074/jbc.RA119.009419 From NLM Medline.

(34) Saccuzzo, E. G.; Martin, M. D.; Hill, K. R.; Ma, M. T.; Ku, Y.; Lieberman, R. L. Calcium dysregulation potentiates wild-type myocilin misfolding: implications for glaucoma pathogenesis. J Biol Inorg Chem 2022, 27 (6), 553–564. DOI: 10.1007/s00775-022-01946-3 From NLM Medline.

(35) Ohashi, R.; Takegoshi, K. Asymmetric (13)C-(13)C polarization transfer under dipolar-assisted rotational resonance in magic-angle spinning NMR. J Chem Phys 2006, 125 (21), 214503. DOI: 10.1063/1.2364503 From NLM PubMed-not-MEDLINE.

