## Supplementary Information for "Evidence for S_331_-G-S-L within the amyloid core of myocilin olfactomedin domain fibrils based on low-resolution 3D solid-state NMR spectra"

**Corresponding Authors**

A

LKESPSGYLRSGEGDTGCGELVWVGEPLTLR
TAETITGKYGVWMRDPKPTYPYTQETTWRID
TVGTDVRQVFEYDLISQFMQGYPSKVHILPR
PLESTGAVVYSGSLYFQGAESRTVIRYELNT
ETVKAEKEIPGAGYHGQFPYSWGGYTDIDLA
VDEAGLWVIYSTDEAKGAIVLSKLNPENLEL
EQTWETNIRKQSVANAFIICGTLYTVSSYTS
ADATVNFAYDTGTGISKTLTIPFKNRYKYSS
MIDYNPLEKKLFAWDNLNMVTYDIKLSKM

B

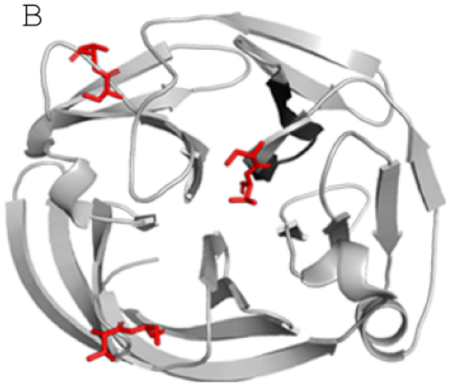

**Supplemental Figure S1. Location of crosspeaks identified from 2D-DARR in the OLF sequence and structure.** 2D-DARR experiments are non-directional, so the identified "S-T" crosspeaks could also be attributed to "T-S" connections. In order to determine which region(s) of OLF may be responsible for forming the fibril core, the OLF sequence and structure were investigated. (A) All possible ST/TS connections were mapped to the OLF sequence in red. P1 (GAVVYSGSLYFQ)is underlined. (B) One of the occurrences of "ST" in the OLF sequence occurs immediately before the P1-peptide sequence (black).

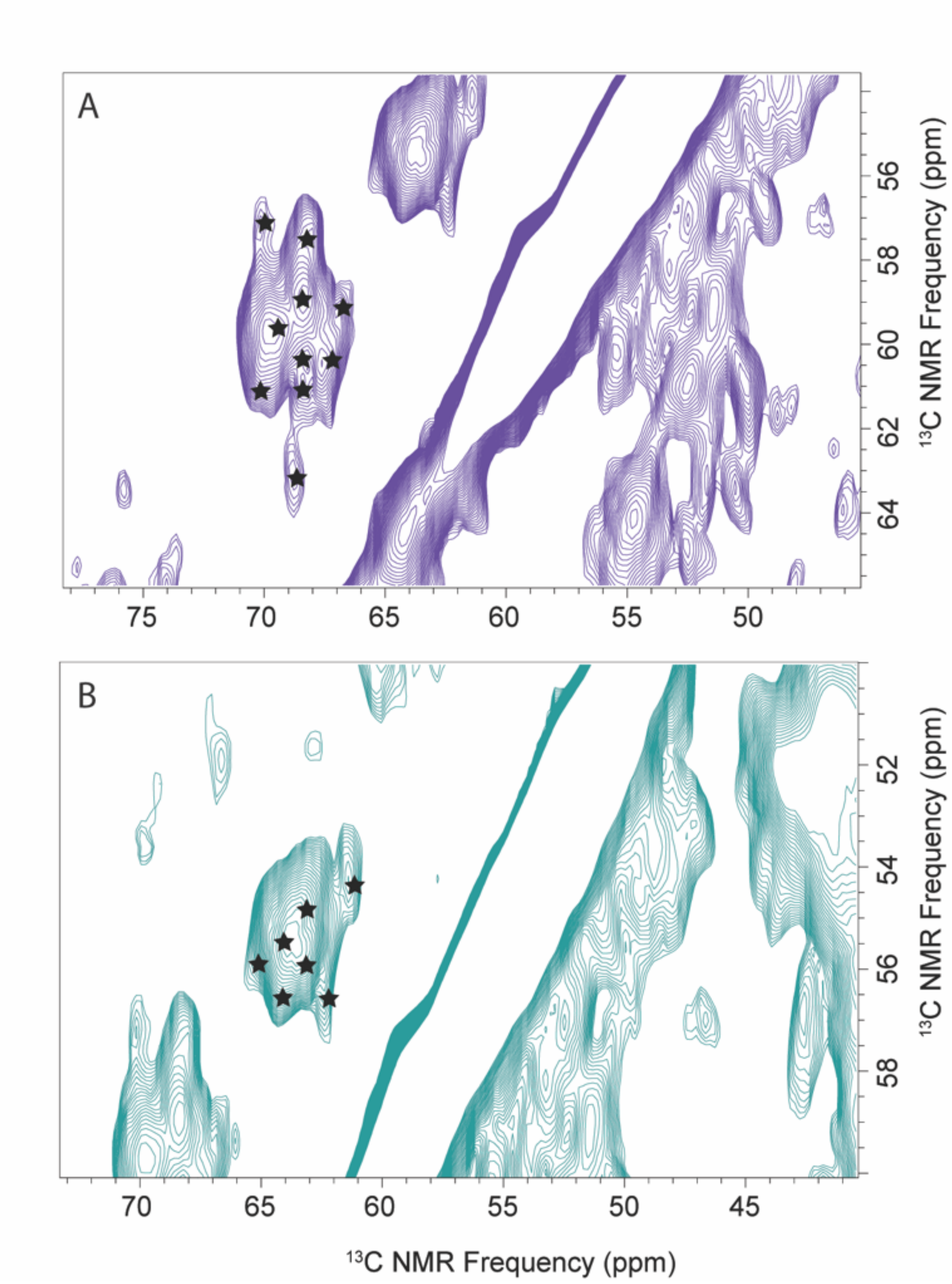


**Supplemental Figure S2. 2D projections of the NCACX spectra are used to count the number of (A) threonine or (B) serine residues in the OLF fibril core.** Out of 20 total serine and 28 total threonine residues in the OLF sequence, only 7 serine and 10 threonine residues were detected in the 3D data from Gaussian peak fitting. This corresponds to 35% of serines and 36% of threonines, suggesting ~36% of the 277 amino acids in OLF (~ 98 residues) are involved in forming the fibril core. However, multiple peaks from different serine residues could overlap and reduce the estimated number of residues, meaning < 98 residues may form the fibril core.

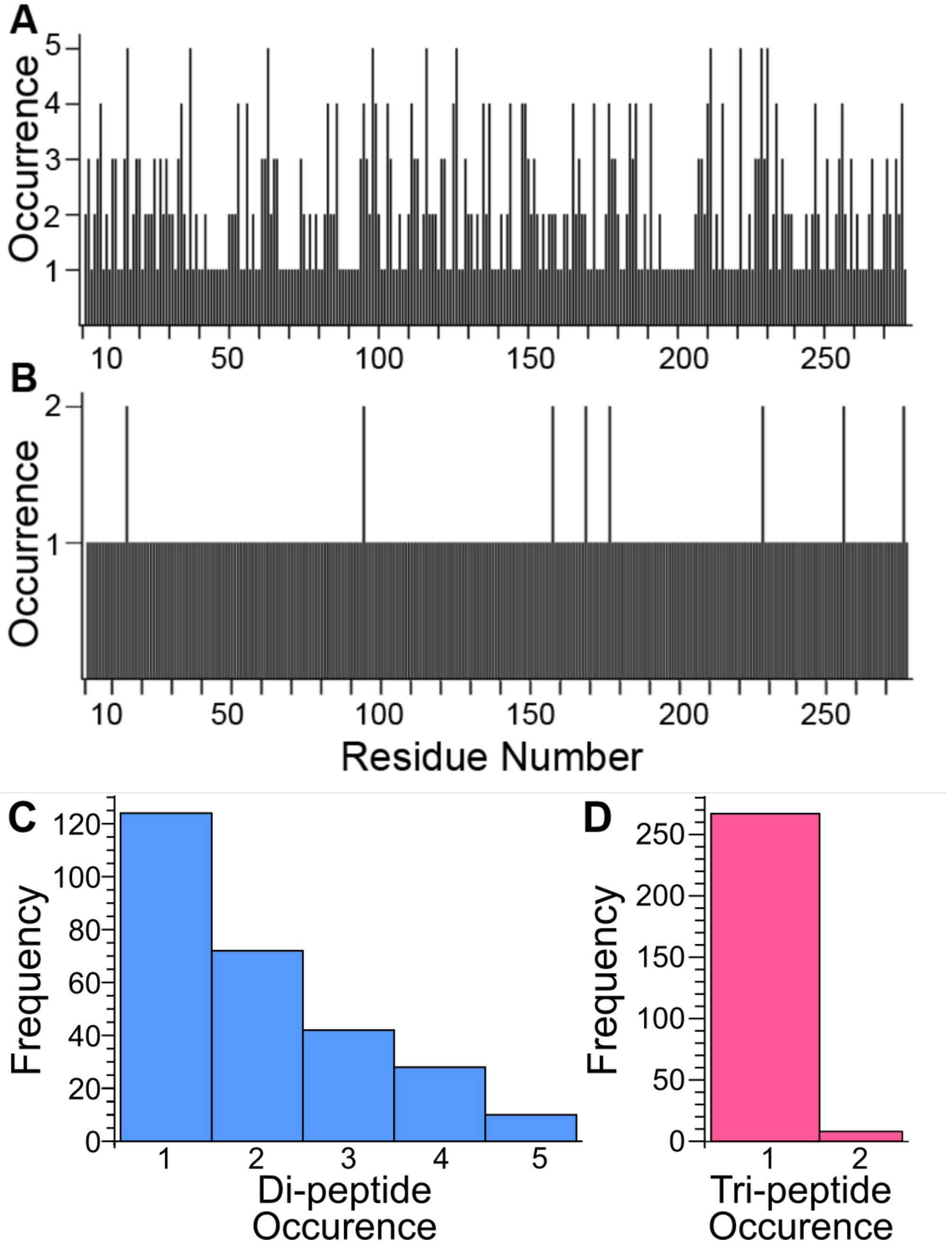


**Supplemental Figure S3. Quantification of unique di and tri-peptides in OLF.** The OLF sequence was analyzed and the occurrence of each set of (A) di- and (B) tri-peptides was quantified, revealing a high number of unique two and three residue sequences that only occur once within OLF. Histograms were made to summarize the frequency of (C) di- and (D) tri-peptide occurrences. The percentage of unique di-peptides in the OLF sequence is 45%. The percentage of unique tri-peptides is 96%.

**Supplemental Table S1. Approximate number of Ser, Gly and Thr residues detected in NCACX/NCOCX spectra.**

| Residue | # in OLF sequence | # detected in NCACX/NCOCX | % |
|---|---|---|---|
| Ser | 20 | ~7 | 35% |
| Thr | 28 | ~10 | 36% |

**Supplemental Table S2. Chemical shifts for all assigned atoms.**

| Residue | N | Cα | Cβ | C' |
|---|---|---|---|---|
| S331 | ? | 55.9 ppm | 64.2 ppm | 171.1 ppm |
| G332 | 115.9 ppm | 44.5 ppm | - | 169.2 ppm |
| S333 | 121.0 ppm | 55.4 ppm | 63.4 ppm | 171.0 ppm |
| L334 | 123.2 ppm | ~ 52.5 ppm | ~ 41.5 ppm | ? |
